# Estimating how Site-Level Differences in Acoustic Environments Affect Species Detection by Machine Learning Models

**DOI:** 10.1101/2025.08.31.673350

**Authors:** Ruari Marshall-Hawkes, Simon Gillings, Mark W. Wilson, Anthony S. Wetherhill, Lynn V. Dicks, Adham Ashton-Butt

## Abstract

1. Machine learning models are vital in species detection from acoustic data, due to the large amount of recordings produced, especially in passive acoustic monitoring studies. However, the acoustic environment of a site can significantly impact the detectability of species using such models, potentially leading to biased estimates of species abundance and distribution. This impact is poorly understood, and current manual validation methods for assessing acoustic detectability are intractable when working with many species, across multiple locations and datasets, due to the intensive time requirements to manually identify false negatives.
2. We introduce a new method, Noise-Augmented Detectability Estimation (NADE), to estimate the impact of site-specific environmental noise on the acoustic detectability of different species. We use vocalisations augmented with noise from the sites themselves to alleviate the need for manual validation of detectability. Our approach produces site- and species-specific estimates of the complex relationship between background noise, vocalisation amplitude, and acoustic detectability when using a machine learning detection model. We evaluate this method using BirdNET on six bird species, in the UK, demonstrating its potential for broader application in acoustic ecology.
3. Our results show that there are large differences in the ability of a machine learning model to detect and classify vocalisations across different sites and species. For example: the detectability of loud vocalisations can be 90% at one site and 10% at another; the detectability at a site can be halved during the course of a survey; and that the amplitude of the background noise alone is insufficient to correctly estimate the detectability of a vocalisation. Detectability at a site can also vary considerably by species: a site can have the relatively high detectability for one species but poorer detectability for another.
4. By providing a quantitative measure of how site-specific background noise affects acoustic detectability, NADE could be used to improve the accuracy of species abundance estimates derived from passive acoustic monitoring, helping to identify sites and times where the detectability is likely to be lower. This approach provides a significant step towards making acoustic detectability estimation more feasible, allowing more robust estimates of species abundance and distribution from acoustic data.

## Introduction

Widespread declines in biodiversity are well established, with many species experiencing significant reductions in range and population size (Inger et al. 2015; Baker et al. 2019; Burns et al. 2021; Sánchez-Bayo & Wyckhuys 2021; Wernberg et al. 2024). Understanding the conservation status of a species relies on data about geographic range and populations. Occupancy and abundance models provide insight into species distributions and range dynamics (Devarajan et al. 2020), community composition, population trends, and associations with environmental variables, all of which can inform targeted conservation (Waldock et al. 2022). Long-term data are especially valuable, as extended time series greatly increase statistical power to detect population trends (White 2019), enabling conservation action before serious declines occur. However, global monitoring efforts remain spatially biased (Moussy et al. 2022), highlighting the urgent need to scale up monitoring programs across ecosystems.

Passive acoustic monitoring (PAM) provides a potential solution to monitor biodiversity for sound producing taxa, relatively cheaply and over large scales (Ross et al. 2023), enabling continuous, long-term sampling across broad spatial extents. PAM data can be used to model occupancy through presenceabsence (Wood & Peery 2022; Cole et al. 2022; Brunk et al. 2023; Kelly et al. 2023; Bielski et al. 2024) and relative abundance via vocal activity indices (Sebastián-González et al. 2018; Pérez-Granados et al. 2019; Pérez-Granados & Traba 2021; Hutschenreiter et al. 2024; Jarrett & Willis 2024). Getting robust occupancy or activity estimates from PAM typically requires extensive data collection across multiple sites and time periods to achieve good survey coverage, especially when aiming to survey multiple species and taxa (Lindner et al. 2025).

Given the scale of PAM data, deep learning, often in the form of convolutional neural networks, underpins many methods for estimating species occupancy and abundance (Stowell 2022; Xie et al. 2023). Deep learning models enable automated detection across diverse taxa including bats, marine mammals, anurans, insects, fish, and birds (Stowell 2022). For detecting bird vocalisations in PAM data, BirdNET (Kahl et al. 2021) and Perch (Ghani et al. 2023) are commonly used. BirdNET takes 3-second audio segments and outputs predicted confidence (0-1) for each of >6,000 trained bird species. BirdNET has been used extensively in occupancy modelling (Cole et al. 2022; Kelly et al. 2023; Brunk et al. 2023; Bielski et al. 2024), and Perch has recently been used in abundance estimation (Navine et al. 2024), with a key insight being the explicit estimation of detectability. However, detectability is often neglected in PAM surveys due to the difficulty of measuring it.

Detectability is a well-known issue in human-based observational surveys of wildlife (Kéry & Schmidt 2008; Guillera-Arroita 2017; Burns et al. 2019; Bennett et al. 2024). If detectability varies (e.g. between sites or species), it needs to be measured to make meaningful insights into abundance changes (Callaghan et al. 2024). In physical surveys, detectability can vary with distance (Buckland et al. 2015), habitat (Gu & Swihart 2004), observer effects (Schmidt et al. 2023), and seasonal factors (Burns et al. 2019). Detectability may also pose a challenge in passive acoustic monitoring (PAM), where variation in detectability could disproportionately affect vocal activity metrics, potentially biasing comparisons across sites, times, or conditions if not accounted for.

Acoustic detectability is influenced by multiple factors (Simons et al. 2009) including distance to the recorder (Pérez-Granados 2023; Winiarska et al. 2024), habitat effects on sound attenuation (Forrest 1994; Darras et al. 2016; Abrahams & Geary 2020; Haupert et al. 2023), environmental conditions (Thomas et al. 2020; Serrurier et al. 2024), species traits (Johnston et al. 2014), and background noise (Haupert et al. 2023). While playback experiments can measure many of these factors (Darras et al. 2016; Thomas et al. 2020; Shaw et al. 2022; Haupert et al. 2023; Pérez-Granados 2023; Hutschenreiter et al. 2023; Lebeuf-Taylor et al. 2024; Winiarska et al. 2024), they are limited to specific times and locations, making large-scale detectability assessment challenging.

Measuring detectability from machine learning models requires evaluating model recall, which is the proportion of true vocalisations correctly detected by a model. Measuring recall is time-consuming, however, as it requires manual validation of false negatives across the full dataset, which becomes prohibitive for multiple species and sites (Sethi et al. 2023). Haupert et al. 2023 demonstrated that background noise is the main environmental feature influencing detection distance. While deep learning models are trained with data augmentation to be invariant to background noise (Stowell et al. 2019; Kahl et al. 2021), the diverse and site-specific nature of background noise (Slabbekoorn 2004; Luther & Gentry 2013) that varies temporally (Haupert et al. 2023) makes comprehensive training coverage impossible. Machine learning models may miss detections through various mechanisms: complete masking by noise, prioritisation of simultaneous vocalisations due to training bias, or masking of algorithm-critical vocalisation segments. Critically, if background noise correlates with ecological variables of interest, detectability variation may bias inference by conflating true ecological patterns with detection artifacts.

In this research, we aimed to quantify how acoustic detectability varied across sites and species during a passive acoustic monitoring survey. We developed a method to quantify the impact of site-specific environmental noise on acoustic detectability of vocalising species, which we call Noise-Augmented Detectability Estimation (NADE). NADE is applicable to any detection model that outputs a confidence score and where background noise influences detectability. We focused on how site-specific background noise affects detectability and how the relationship between vocalisation amplitude and detectability varies between sites and species. We hypothesised that both background noise amplitude and vocalisation amplitude are strongly related to a machine learning model’s to detect and classify vocalisations, and that these factors vary spatio-temporally and between species. We tested NADE using BirdNET detections of six bird species across 28 sites surveyed for 1.5 months each. Our approach makes acoustic detectability estimation more tractable for large-scale biodiversity monitoring by reducing manual validation requirements and potentially improving confidence in PAM-derived abundance estimates.

## Materials and Methods

### Study Sites and Species

We used acoustic data from a survey of 28 sites spread across Scotland, from 7°W to 2°W, 55°N to 59°N, with a total dataset size of 2,060 hours of audio. Each audio recorder was set to record on a schedule of 1 minute on followed by 14 minutes off. The recorders were deployed continuously for an average of 46 days (range of 28 to 68 days) between 17^th^ April and 7^th^ July 2023. This period encompassed or overlapped with the breeding season for most of the bird species found at these sites.

This coincided with in-person monitoring at the same sites using the Breeding Bird Survey line transect method (Massimino et al. 2025). Most audio recorders were deployed during the first in-person survey, and were collected during the second in-person survey. The recorders used were Song Meter Micro with a sample rate of 22050Hz and a gain of 18dB.

We developed and tested NADE on six common UK bird species (Tree Pipit (*Anthus trivialis*), Meadow Pipit (*Anthus pratensis*), Goldcrest (*Regulus regulus*), European Greenfinch (*Chloris chloris*), Common Cuckoo (*Cuculus canorus*), and European Golden-Plover (*Pluvialis apricaria*)). These species were chosen because they were present and vocalising within many of the study sites, have distinct vocalisations, and are spread across a range of vocalisation frequency ranges. NADE can be applied to any species that vocalises and is detectable by an AI model.

### Measuring the impact of site-specific noise on acoustic detectability

To estimate the impact of site-specific noise and detected vocalisation amplitude on detectability we simulated detections by augmenting foreground vocalisations of known amplitude with site-specific background noise and measuring the resulting model confidence scores. For our detection and classification model, we used BirdNET^1^.

1. **Initial BirdNET predictions:** We first ran BirdNET on the entire audio dataset, detecting and classifying the species present within every 3-second chunk of audio without overlap, and recorded the confidence score associated with each classification.
2. **Foreground audio clip selection:** Then, we took predictions with confidence *≥* 0.98 for each species. These high-confidence clips likely represent good quality, low-noise vocalisations that we aim to detect consistently. Importantly, we selected foreground audio from the dataset itself, rather than external sources like Xeno-canto (Xeno-canto Foundation 2025), to ensure the frequency response of the foreground audio clips is consistent with true vocalisations that occurred at the study sites. We then sampled 15 vocalisations across the range of different vocalisations that the species makes and validated them by looking at and listening to the associated 1-minute spectrogram to confirm they were the correct species. We measured the minimum and maximum vocalisation frequency of each vocalisation, considering only the fundamental frequency as in Leseberg et al. 2022, and computed the amplitude within this range across the duration of the vocalisation.
3. **Background audio clip selection**: For each site, we randomly sampled 1000 3-second audio clips from across the survey period between 4am and 9pm, aligning with typical daylight hours for the study sites. These form our site-specific noise profiles. The same background audio clips are used for each species to allow for easier comparison. For each background clip we measured the amplitude within the frequency range of the vocalising species. We measured the amplitude of an audio clip using the root mean square, and these amplitudes are reported in decibels relative to full scale (dBFS), where 0dBFS represents the maximum possible amplitude the recorder can capture, and smaller values indicate proportionally quieter signals. The self-noise of the recorders used is approximately -70dB.
4. **Augmenting audio clips with background noise:** For every background / foreground clip combination, we randomly sampled a vocalisation amplitude value uniformly in the range [-80dBFS, -20dBFS]. Then, the foreground clip in that combination had its original amplitude adjusted to match the sampled vocalisation amplitude by adding or removing gain (i.e. adjusting the signal’s amplitude by increasing or decreasing the volume). This approximated the range of detected vocalisation amplitudes within the dataset from quiet to loud. The foreground clip was then augmented with each background clip by stacking the audio files (i.e. summing the waveforms). If the background clip contained a detection of the foreground clip species with confidence > 0.1, then this background/foreground combination was skipped.
5. **BirdNET predictions of augmented audio clips:** For every site, we ran BirdNET on the augmented audio clips and recorded the confidence. Then, we determined whether each augmented audio clip was ‘detected’ based on a confidence score threshold. We then measured the impact of background noise on detectability as the proportion of augmented audio clips that were detected.

Our workflow results in 15,000 augmented audio clips per site for a single species. For each augmented audio clip we know the vocalisation amplitude, the background noise amplitude, the site and date/time from which the background noise sample was taken, and the prediction confidence score. This allows us to model the relationship between signal, noise, and detectability across a range of vocalisation types, background noise profiles, and detected vocalisation amplitudes. This process is repeated for each site and for each species, allowing us to see how these influence detectability. The full workflow is in (Fig. 1).

**Figure 1.**
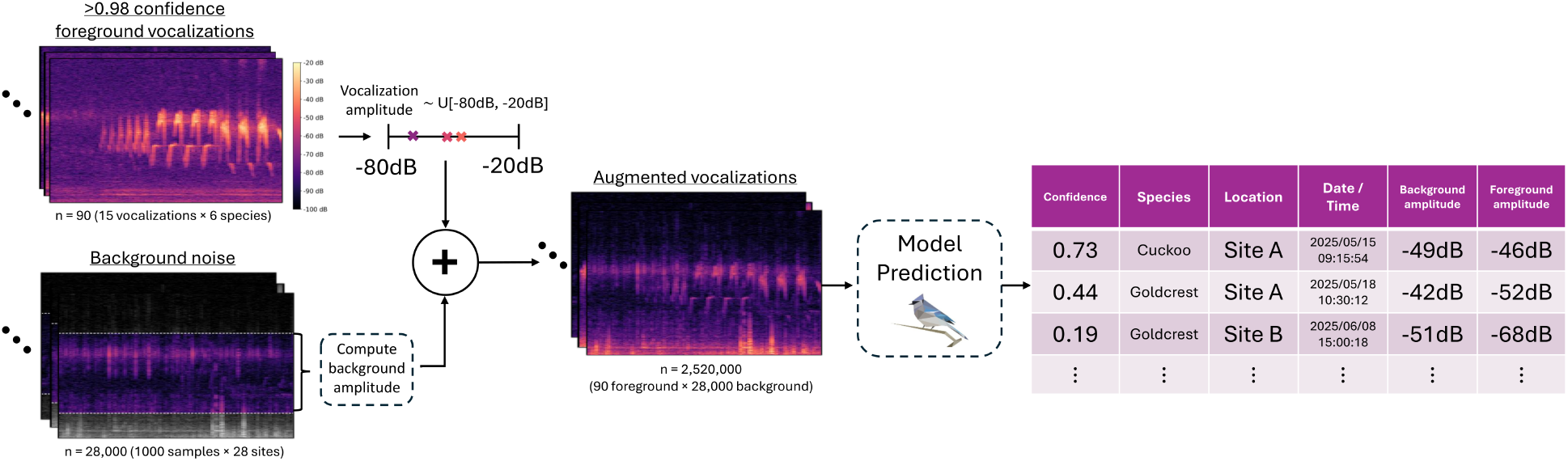
An overview of NADE, used to estimate the impact of background noise on acoustic detectability. High confidence foreground vocalisations are sampled and validated, and their amplitude is sampled uniformly from the range (-80dBFS, -20dBFS). These are then augmented with background noise samples. BirdNET is run on these augmented clips, generating an output table with the confidence, site name, background date/time, foreground amplitude, and background amplitude.

To investigate the impact of the type of background noise on detectability, we labeled background noise samples containing only bird or wind noise. We then generated white noise across a range of amplitudes. This lead to 1,560 background noise samples in each of the three categories. Finally, we apply the same workflow as in the other experiments to measure the impact of background noise type on detectability whilst controlling for the background noise amplitude.

### Statistical Modelling

We modeled the probability of detection as a binary variable detected, whose value depended on whether on not the vocalisation was detected. We used a generalized linear mixed model (GLMM) with a binomial error distribution and a logit link function (Bolker et al. 2009). We use the R package glmmTMB (Brooks et al. 2017). In all models we accounted for pseudoreplication by adding two nested random effects: (1 | species:vocalisation), which accounts for the same 15 vocalisations per species being used for all sites, and (1 | location:background), which accounts for the same 1,000 background noise samples per site being used for all species. In all models, Site E was the reference location and Meadow Pipit was the reference species, such that the intercept represents the predicted detectability when the site was E and the species was Meadow Pipit. The final number of datapoints used is 2,485,725: 6 species × 15 vocalisations × 28 locations × 1,000 background noise samples, exlcuding the combinations where the background noise contained a detection of the foreground clip species with confidence > 0.1.

### Modelling the impact of site-specific noise

To test if the location is an important factor in the detectability of high amplitude (-30dB to -20dB) vocalisations, we use a likelihood ratio test on the following two models:

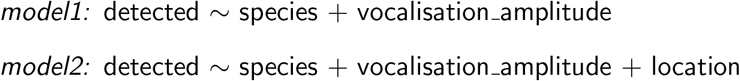

To test whether detectability differed significantly between sites for each species, we used estimated marginal means from the fitted model and performed Tukey-adjusted pairwise comparisons of site effects.

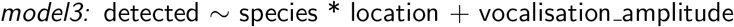

To test how similarly each species ranked sites by detectability of high-amplitude vocalisations, we first used estimated marginal means from *model3* to obtain predicted detectability for each site and species. Within each species, sites were ranked in order of predicted detectability. We then computed a consensus site ranking as the mean rank across all species and calculated the Spearman rank correlation between each species’ site rankings and this consensus ranking.

### Modelling the impact of vocalisation amplitude

To test if vocalisation amplitude is an important factor in detectability, we use a likelihood ratio test on the following two models:

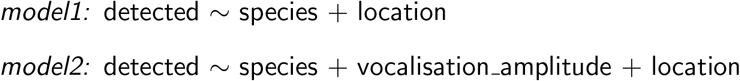

To test if the impact of vocalisation amplitude on detectabiltiy varies by location and by species, we assessed the significance and effect sizes of the interaction terms in *model3* relative to the main effect of vocalisation amplitude:

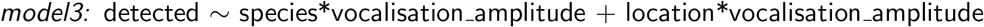

### Modelling the impact of background noise amplitude

To test if background noise amplitude is an important factor in detectability, we use a likelihood ratio test on the following two models:

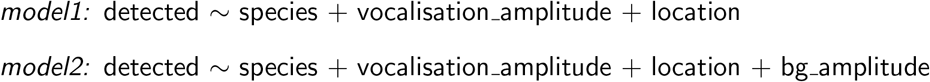

To test if the impact of background noise amplitude on detectabiltiy varies by location and by species, we assessed the significance and effect sizes of the interaction terms in *model3* relative to the main effect of background noise amplitude:

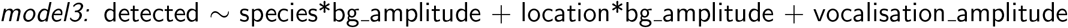

To test whether the impact of background noise amplitude on detectability differs between low- and high-amplitude vocalisations, we included the factor amp group (*high* if -30 < vocalisation amplitude < -20 and *low* if -50 < vocalisation amplitude < -40) and its interactions with background noise amplitude in *model4* below. The significance of the amp group:bg amplitude interactions was used to assess whether the effect of background noise on detectability varied between amplitude groups.

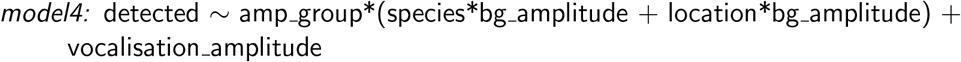

To assess whether lower-amplitude vocalisations showed greater variability in their responses to background noise, we estimated the slopes of detectability vs. background noise amplitude for each species and location within each amplitude group (using marginal trends from *model4* ). We then compared the range and dispersion of these slopes between the low- and high-amplitude groups.

To assess if the type of background noise impacts detectability, we assessed the significance and effect sized of the noise type (bird, wind, or white noise) in *model5* relative to the main effect of background noise amplitude, with white noise as the reference noise type:

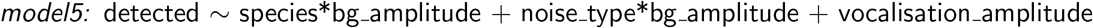

## Results

### Site-specific noise leads to different detectability at each site

By augmenting the vocalisations with background noise from sites we are able to isolate the impact of site-specific noise on species detectability. The distribution of confidence values for high amplitude vocalisations between -30dB and -20dB is given in Fig. 2 for five of the sites across the survey period. We selected five sites of out the 28, A through E, to illustrate the range of possible outcomes and highlight them among the full dataset in Figures 2-5. The results suggest that there are differences between sites in their distribution of confidence values and, therefore, their detectability, and that these distributions vary across species. Even for the loudest, and therefore easiest-to-detect vocalisations, background noise had an impact on detectability.

**Figure 2.**
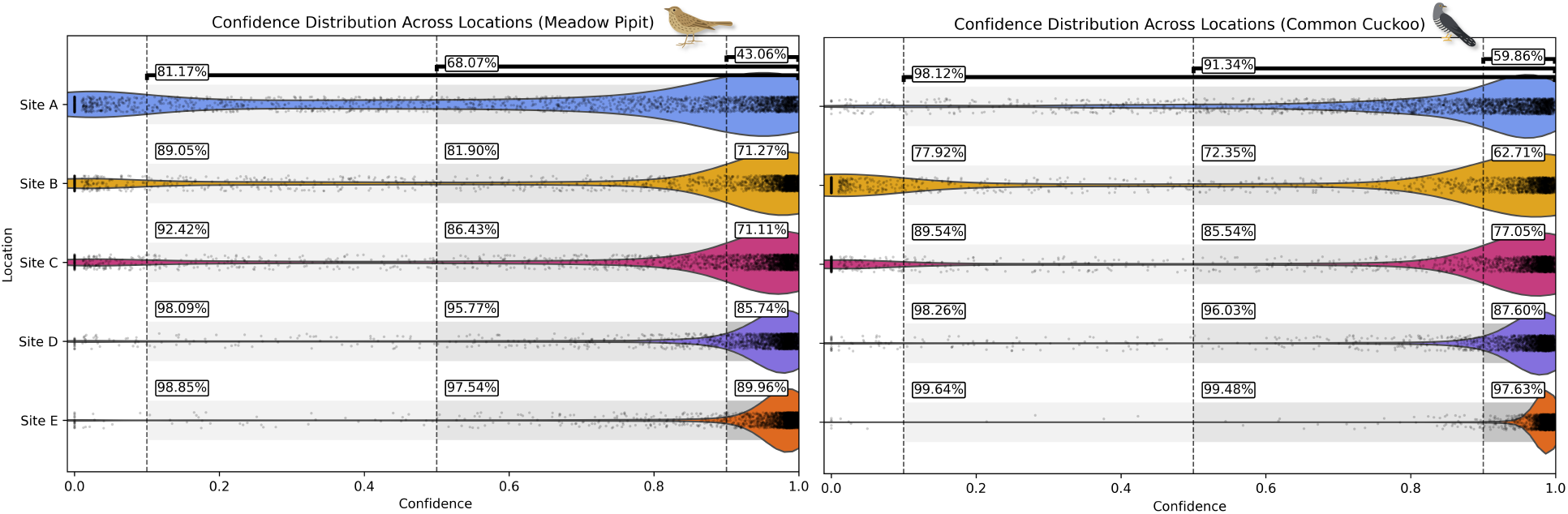
The different distributions of BirdNET prediction confidence values for the augmented foreground vocalisations for five sites (A – E) and two species (left: Meadow Pipit, right: Common Cuckoo), using the same set of vocalisations at each site with only the background noise varying. Each scattered point represents a single vocalisation-background audio clip combination for vocalisation amplitudes between -30dBFS and -20dBFS, and the density of these points is given by the violin plot. For each site, the percentage of augmented clips whose confidence score is above 0.9, 0.5, and 0.1, is shown to right of the respective confidence score threshold. A version of this plot with all species is given in Fig. S2.

The detectability of augmented vocalisations differed across sites for each of the six species (*χ*^2^ = 4, 599, df = 27, *p* < 0.0001) at all confidence tested. At a 0.9 confidence threshold, detectability ranged from 0.43 to 0.99 across sites, with mean detectability values by species of Meadow Pipit (0.83), Common Cuckoo (0.85), Tree Pipit (0.89), European Greenfinch (0.91), Goldcrest (0.91), and European Golden-Plover (0.92). 72–82% of site comparisons yielded significant differences (*p* < 0.0001). For each species, we ranked sites by their detectability. The rankings of sites were broadly consistent across five of the six species (Spearman correlation with the consensus > 0.95), while Common Cuckoo had much lower correlation with the consensus (Spearman *ρ* = 0.64), implying site-specific noise impacted Common Cuckoo differently to the other species. Using a conservative threshold of 0.9, for -30dB to -20dB Meadow Pipit vocalisations, 90% were detected at Site E, compared to just 43.1% at Site A (Fig. 2) – that is, the recall of an individual vocalisation was 2× higher at Site E. With a low confidence threshold of 0.1, vocalisations were hardest to detect at Site A, with only 81.2% of vocalisations detected. In contrast, Common Cuckoo vocalisations were relatively easy to detect at Site A, with 98.1% of vocalisations detected.

### Vocalisation amplitude is strongly related to detectability

The amplitude of a vocalisation had a large impact on its detectability (*χ*^2^ = 1, 980, 775, df = 1, *p* < 0.0001). This relationship between vocalisation amplitude (*β*_vocalisation_ = 0.294) and detetability varied by site (24/27 with *p* < 0.05, *β*_vocalisation:site_ between -0.110 and 0.039 with median -0.012) and by species (all *p* < 0.001 *β*_vocalisation:species_ between 0.019 and 0.033 with median 0.022) (Fig. 3). For every species there is a clear difference in the relationship between vocalisation amplitude and detectability at a 0.9 confidence threshold between sites, with the same vocalisation amplitude leading to very different detectability across sites. For European Greenfinch, -30dB vocalisations at Site A are detected 40% of the time, whereas at Site B they are detected 77% of the time. Critically, it is not until the vocalisation at Site B is at -60dB when the detectability is the same as the -30dB vocalisation at Site A. This suggests that, for any given detected European Greenfinch vocalisation at around -30dB at Site A, if the same vocalisation occurs at Site B it will have almost the same detectability, even if it’s amplitude is 32× lower. When vocalisations are lower amplitude, for example between -50dB and -40dB, the differences in detectability between sites is exacerbated compared to high amplitude vocalisations between -30dB and -20dB. For example, a -50dB Greenfinch vocalisation had just 2% at Site A, but 62% at Site B. For all species, very low amplitude vocalisations below -70dB (just above the self-noise floor of the microphone) lead to detectability close to zero at every site.

**Figure 3.**
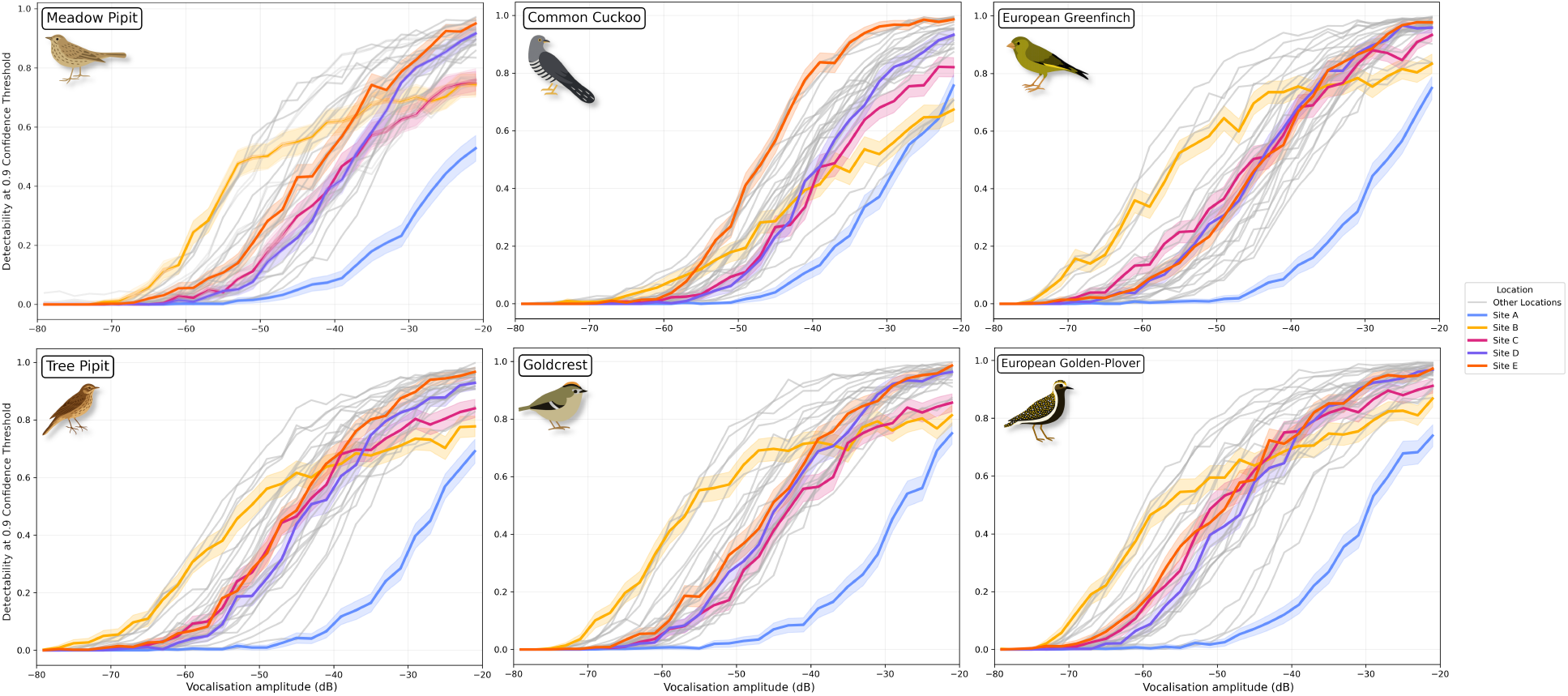
The relationship between the detectability of augmented audio clips at a 0.9 confidence threshold and vocalisation amplitude for the six study species at each of the sites. Each line shows a different site, with the detectability computed for 2dB bins of vocalisation amplitude. 95% confidence intervals are given by the shaded regions. Five of the 28 sites (A – E) are highlighted in colour.

For most species, certain sites consistently showed higher detectability, with detectability increasing at a similar rate as amplitude increased. This pattern was not consistent across all species, however. For Common Cuckoo, Site E had the highest detectability across the amplitude range (detectability rank 1/28), whereas for other species Site E was typically average (median detectability rank 13/28). In contrast, Site B had particularly low detectability for Common Cuckoo (detectability rank 25/28) in comparison to other species (median detectability rank 7/28). Notably, no site achieved more than 50% detectability for the low amplitude -50 dB Common Cuckoo vocalisations, whereas a median of 5/28 sites did for other species, and 13/28 sites did for European Golden-Plover.

### Temporal changes in site-specific noise lead to differences in detectability across the duration of a survey

Detectability varied considerably throughout the survey period (Fig. 4) due to changes in background noise over time. The amount of variation within each site ranged from 0.05 to 0.67, with a median range of 0.20 across all sites. For Meadow Pipit, at Site B, detectability fell from 94% to 31% within 3 weeks, then rose again throughout the rest of the 64 days the audio recorder was deployed. At Site C, the detectability was more constant, though it increased from 50% up to 72% towards the end of the survey. Both Site A and E slowly decreased detectability throughout the survey period, whereas Site D remained mostly constant with only small variations. On average, Meadow Pipit detectability varied by 26% (95% CI: *±* 6%) within a site due to impacts of background noise alone. These temporal variations in detectability were not consistent across species. For example, at Site E, the detectability of Meadow Pipit decreased across the duration of the survey, whereas for Common Cuckoo it increased. Amongst the non-highlighted sites, detectability also showed a large degree of temporal variation, with some sites decreasing over time, some increasing, and some oscillating between periods of high and low detectability.

**Figure 4.**
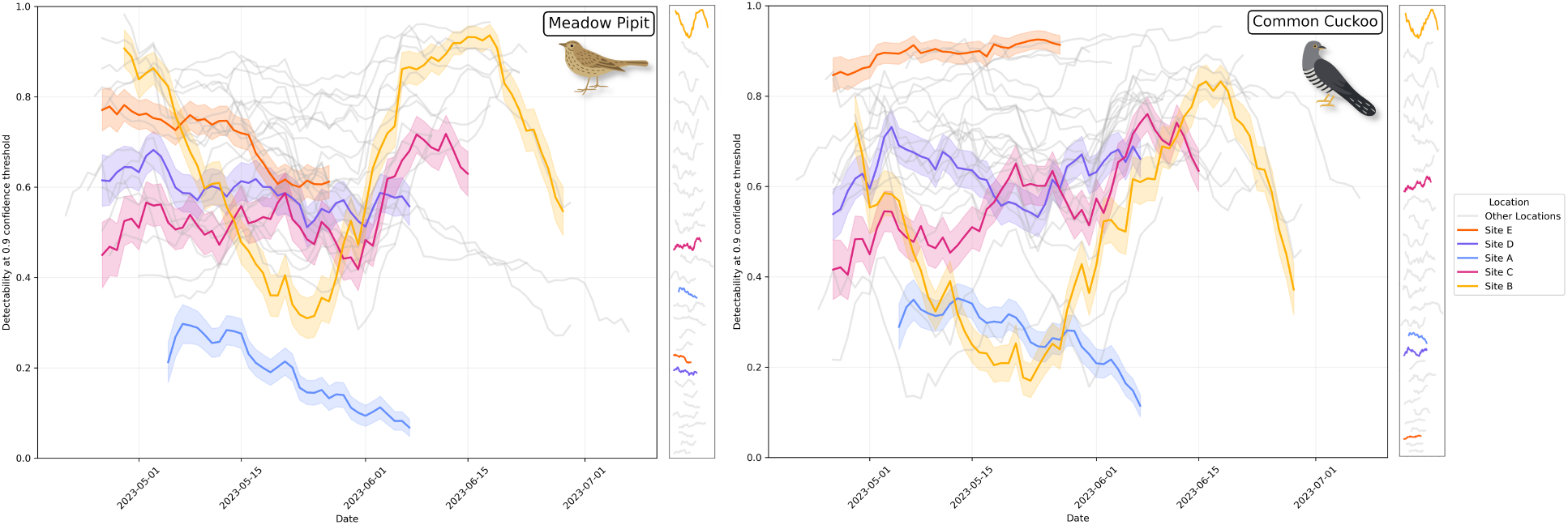
The detectability of a vocalisation at a 0.9 confidence threshold, computed across 10-day bins, for Meadow Pipit and Common Cuckoo vocalisations with amplitudes between -40dB and -30dB. Beside each graph the lines are shown from highest (top) to lowest (bottom) range in detectability over the course of the survey. Five of the 28 sites (A – E) are highlighted in colour. A version of this plot with all species is given in Fig. S3

### Amplitude of background noise at a site is not sufficient to estimate detectability

Background noise had a large impact on the detectability (*χ*^2^ = 103, 911, df = 1, *p* < 0.0001). However, the relationship between background noise amplitude (*β*_background_ = *−*0.184) and detectability varied greatly across sites (23/27 with *p* < 0.05, *β*_background:site_ between -0.102 and 0.092 with median 0.002) and species (all *p* < 0.001 *β*_vocalisation:species_ between 0.016 and 0.047 with median 0.031). This relationship was also dependent upon the vocalisation amplitude (Fig. 5). For very high amplitude vocalisations between -20 and -30dB (which are rare in a PAM survey), the background noise amplitude appeared to have a strong relationship with detectability. However, at more realistic quieter vocalisation amplitudes between -50 and -40dB, the relationship between background noise amplitude and detectability became much more complex, such that the impact of the amplitude of background noise on detectability varied greatly on a site-by-site basis.

**Figure 5.**
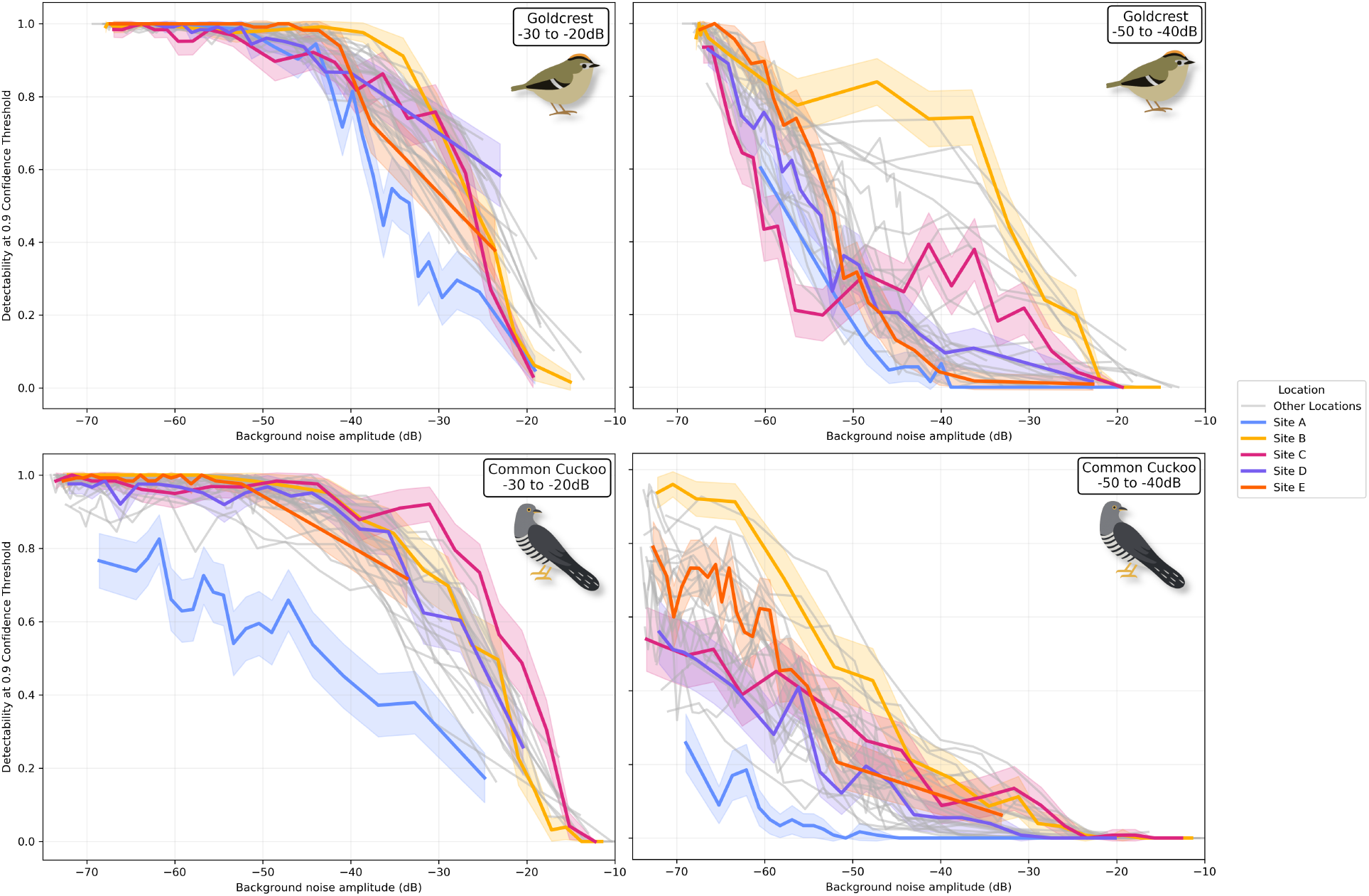
The relationship between background noise amplitude and detectability for Goldcrest and Common Cuckoo. Left column: high amplitude -30dB to -20dB vocalisations. Right column: average amplitude -50dB to -40dB vocalisations. Five of the 28 sites (A – E) are highlighted in colour. A version of this plot with all species is given in Fig. S4.

For both species at high vocalisation amplitudes we saw a small amount of variability in the relationship between background noise amplitude and sites, with Site A being an outlier in that lower amplitude background noise from that site is required to decrease the detectability. At the lower, and more realistic, vocalisation amplitudes of -50dB to -40dB, the relationship between background noise amplitude and detectability became very variable between sites. For example, for Goldcrest, at Site B, -40dB of noise lead to 74% detectability, but at Site A (and 7 of the other sites -40dB of background noise lead to lower than 10% detectability.

We investigated what is causing these differences between sites (see Appendix S1 in Supporting Information). We found that noise containing birds (which were mostly House Sparrow (*passer domesticus*)) impacted detectability intercept more than wind noise when compared to white noise for all species (*β*_intercecpt_ = *−*4.074, *β*_bird_ = 6.103, *β*_wind_ = 4.231, all *p* < 0.001) and the slope (*β*_background_ = *−*0.336, *β*_background:bird_ = 0.201, *β*_background:wind_ = 0.058, all *p* < 0.001).

## Discussion

Our study demonstrates that detectability of vocalisations using machine learning classifiers varies considerably between sites and species, with potentially large effects when modelling relative abundance and occupancy. Noise-Augmented Detectability Estimation (NADE) provides a tractable way of estimating these differences in detectability. We found detectability varies dramatically between sites (e.g., from 62% to 2% for -50dB Greenfinch vocalisations (Fig. 3)), that vocalisation amplitude strongly relates to detectability (though this relationship varies between sites and species), and that detectability fluctuates over time (e.g., from 94% to 31% within 3 weeks for Meadow Pipit at Site A (Fig. 4)). Background noise amplitude alone was not the only factor influencing the detectability of a vocalisation, implying that other factors such as timbre play a role.

NADE rapidly identifies when and where detectability varies in passive acoustic surveys with minimal manual validation – we validated fifteen, 3-second vocalisations per species. Unlike playback experiments (e.g. Haupert et al. 2023) that measure detectability over short timescales, NADE can be used to quantify spatio-temporal variation over the entirety of a PAM survey. Where vocal activity is used as a proxy for species abundance, relative detectability estimates from NADE could standardise vocalisation counts, enabling more robust comparisons between sites and over time.

### Spatio-temporal variation in background noise causes large differences in detectability

We found that background noise caused detectability to vary substantially between sites, such that the same vocalisation was twice as likely detected at one site versus another. Without accounting for differences in detectability in PAM studies, researchers could draw misleading conclusions, particularly for abundance relationships (Callaghan et al. 2024). We found quiet vocalisations at one site can be as detectable as much louder vocalisations elsewhere (up to 30dB difference), implying substantial variation in effective detection range. Using attenuation values from Yip et al. 2017, 30dB differences correspond to sound sources being *∼* 21× further at forest edges or *∼* 12× further in forest interiors.

Detectability also varied temporally within sites. For example, at Site B, over the course of 5 weeks detectability dropped by nearly 70% for Meadow Pipit vocalisations, then subsequently rose. Other sites showed gradual changes or oscillations. These findings partly explain why longer acoustic surveys provide better occupancy and abundance estimates (Cole et al. 2022; Lindner et al. 2025). Though we focused on seasonal variation over the deployment period, we also observed large day-to-day changes, and year-to-year changes are possible in long-term studies with repeated site visits. Failing to account for temporal variation could result in misleading trends. NADE provides fine-scale estimates of the impact of background noise on acoustic detectability, helping to improve the quality of inferences that can be made from PAM datasets without the need for intensive experimentation or manual data collection.

### Background noise amplitude alone is not a reliable predictor of detectability

We showed that background noise amplitude within a species’ vocalisation frequency range is not, by itself, a reliable predictor of detectability, as different sites had different relationships between background noise, amplitude, and detectability. While background amplitude can be a useful first-pass metric to flag periods or sites with very high or low detectability, it did not capture the complex ways that background noise properties – such as their temporal variation and spectral characteristics – affected detectability.

Different noise types affected detectability differently. For example, bird vocalisations or white noise suppressed detectability more than wind noise. We infer that temporal characteristics influence detectability: high amplitude wind noise affecting small vocalisation portions often did not prevent detection, while continuous noise greatly decreased detectability over extended periods, even when the foreground vocalisation was much louder than the background noise.

The acoustic similarity between background noise and vocalisations appears crucial. Background sounds resembling target vocalisations (e.g., other bird calls) likely reduce detectability more than acoustically distinct sounds. Although BirdNET includes noise augmentation (Kahl et al. 2021), it uses only non-bird noise, limiting its robustness in realistic ecological conditions when multiple species vocalise simultaneously. Vocalisation frequency range appears key in determining how site-specific noise affects detectability. For example, Common Cuckoo (400-900Hz) was more affected by low-frequency noise, while Meadow Pipit (2500-8000Hz) was more affected by mid-to-high frequency noise (see Appendix S1). Species with similar frequency ranges showed similar detectability patterns even if their vocalisations are acoustically distinct. For instance, Meadow Pipit and European Greenfinch have vocalisations that are acoustically distinct, but overlap considerably in frequency, and showed similar detectability trends across sites (see Fig. 3).

### Limitations and future work

NADE focuses on background noise and vocalisation amplitude impacts on acoustic detectability, but does not address all relevant factors. We do not estimate site-specific sound attenuation necessary for converting amplitudes to distances, though habitat-specific values can provide reasonable approximations as is demonstrated in Haupert et al. 2023. We only assess site effects on detectability of clear, high-confidence vocalisations and site-specific background noise may have a different relationship with detectability of lower “quality” vocalisations. Bird vocalisations can change in noisy environments (Nemeth et al. 2013; Luther & Gentry 2013), and hardware degradation can also impact detection (Jarrett et al. 2025). Our current approach assumes that recording hardware is consistent across sites and time. While this is likely to be a reasonable assumption for this study, due to the fact that all the recorders were new at the outset, it may not hold for long-term deployments.

Although we identified clear temporal patterns, background noise varies across diel and seasonal timescales (Haupert et al. 2023), and our uniform sampling of 1000 background clips per site may not fully capture this variation. Similarly, 15 vocalisations per species unlikely represent full vocal repertoires, and detectability may differ across vocalisation types. The trade-off between increasing background samples, validating more vocalisations, and computational cost should be systematically evaluated and tailored to the specific context of a study.

NADE does not support direct comparisons of absolute detectability values between species. Differences in selected foreground vocalisations, particularly variation in complexity and initial confidence scores, could create apparent differences reflecting methodological artifacts rather than true differences. More systematic characterisation of call types and amplitude distributions, plus classifier performance assessment for each species, could improve between-species comparisons. However, relative comparisons using NADE remain valid: sites can be ranked by detectability for each species individually, and rankings compared across species. We used this approach to show Common Cuckoo exhibits site-specific detectability patterns differing from other species, providing meaningful insights despite limitations in absolute comparisons.

## Conclusion

This study introduces NADE, a method for estimating the impact of site-specific background noise on species detectability in acoustic monitoring. Our findings demonstrate that background noise significantly affects species detectability, varying across sites and within sites throughout surveys. Factors like other vocalising species and wind noise can dramatically decrease detectability, and the amplitude of noise alone is insufficient to estimate detectability. This approach offers a potential supplementation to manual validation processes, with important implications for improving large-scale passive acoustic monitoring estimates of species abundance through vocalisation rate. By accounting for species-specific spatio-temporal variations in acoustic detectability, the reliability of PAM abundance estimates could be improved, facilitating robust and large-scale ecological monitoring.

## Acknowledgments

We are grateful to Mark Constantine and Linda and Ken Smith for generously funding this work. A huge thanks to the 22 BTO/JNCC/RSPB Breeding Bird Survey volunteers who collected the acoustic dataset we used in this study, and thanks also to the dozens of other volunteers who offered their help. This work was supported by the UKRI Centre for Doctoral Training in Application of Artificial Intelligence to the study of Environmental Risks (reference EP/S022961/1).

## Data Accessibility

The NADE code, along with the vocalisations and background noise clips used in this paper, can be downloaded from GitHub (https://github.com/ruarimh/NADE). NADE is open source under the MIT License.

## Appendix S1

### Detectability is related to background noise type

By inspecting spectrograms, we manually labeled background noise samples as containing either bird noise, or wind noise, or other noise. We also generated white noise samples as a baseline. The wind and bird background noise clips are combined with the foreground clips as in the other results sections.

**Figure S1:**
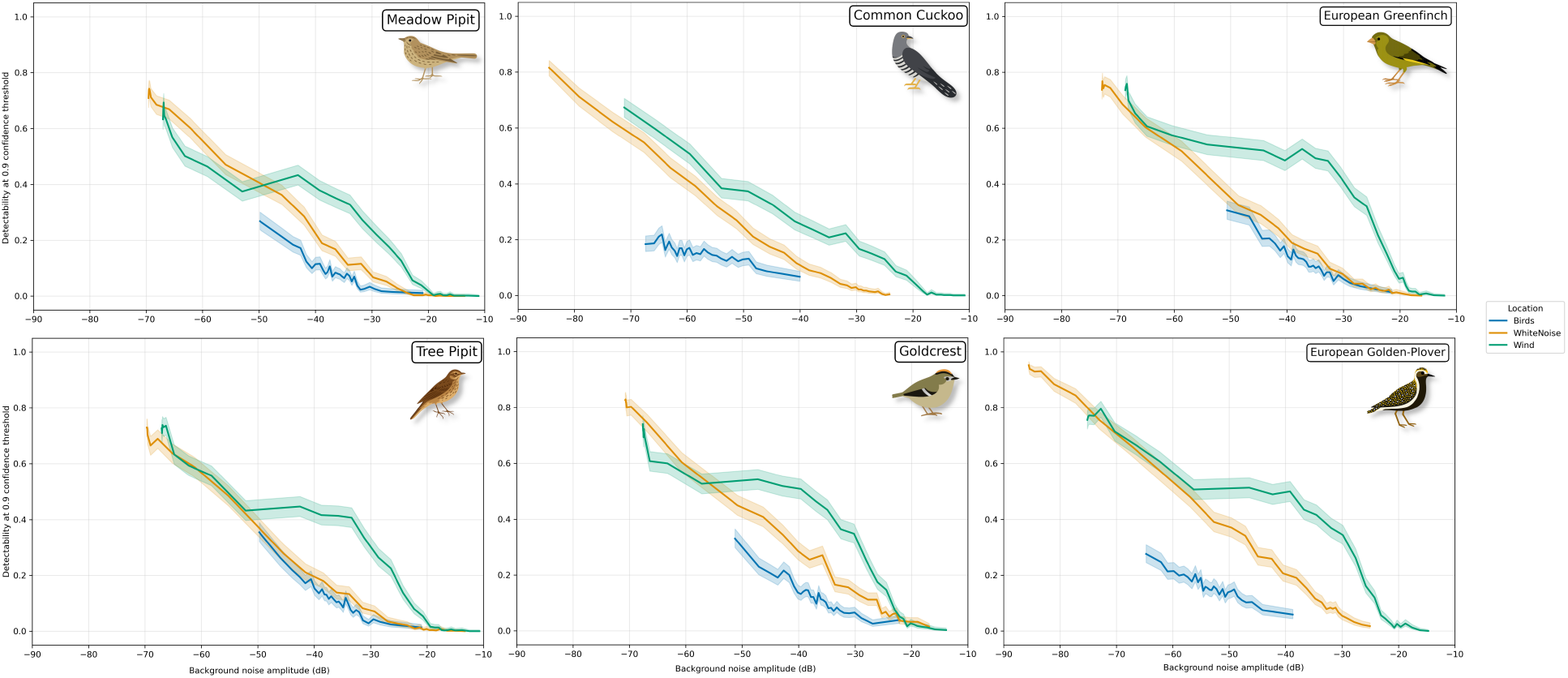
The relationship between background noise amplitude and detectability at a 0.9 confidence threshold for three different types of noise: bird noise, wind noise, and white noise. 95% confidence intervals are given by the shaded regions.

## Appendix S2

### Site-specific noise leads to different detectability at each site

**Figure S2:**
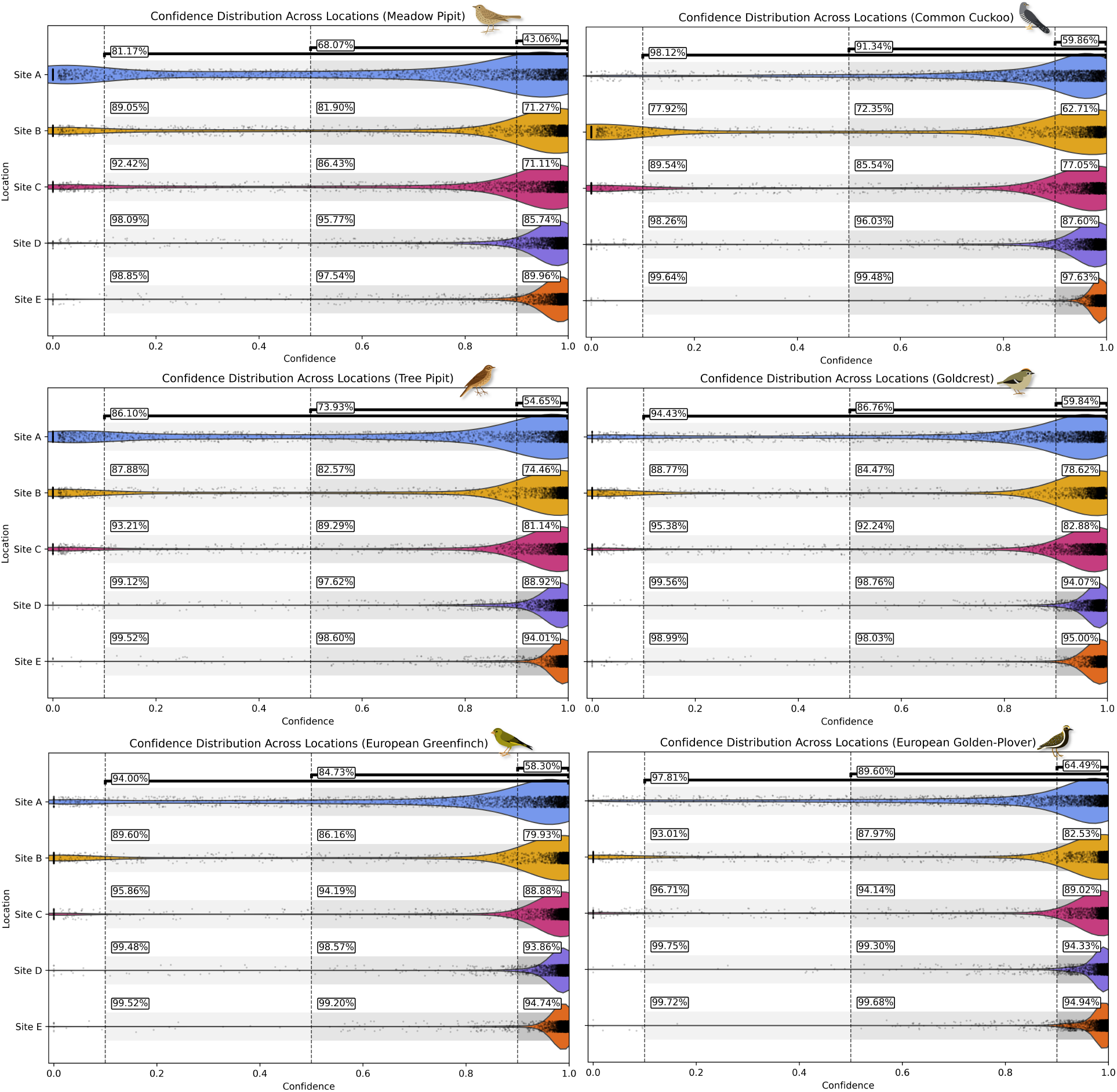
The different distributions of BirdNET prediction confidence values for the augmented foreground vocalisations for five sites (A – E) and all six species, using the same set of vocalisations at each site with only the background noise varying. Each scattered point represents a single vocalisationbackground audio clip combination for vocalisation amplitudes between -30dBFS and -20dBFS, and the density of these points is given by the violin plot. For each site, the percentage of augmented clips whose confidence score is above 0.9, 0.5, and 0.1, is shown to right of the respective confidence score threshold.

#### Temporal changes in site-specific noise lead to differences in detectability across the duration of a survey

**Figure S3:**
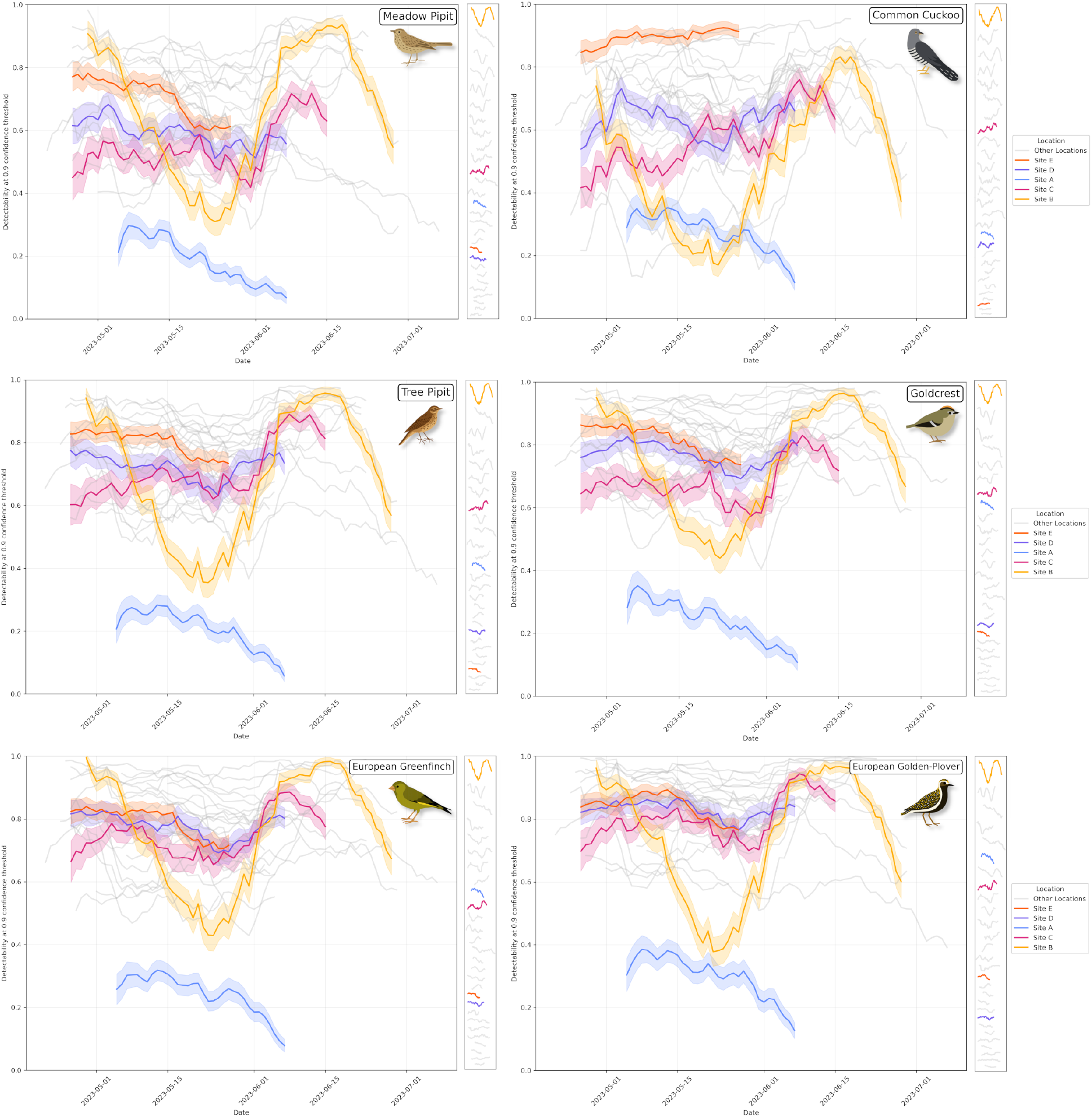
The detectability of a vocalisation at a 0.9 confidence threshold, computed across 10-day bins, for all six species vocalisations with amplitudes between -40dB and -30dB. Beside each graph the lines are shown from highest (top) to lowest (bottom) range in detectability over the course of the survey. Five of the 28 sites (A – E) are highlighted in colour.

#### Amplitude of background noise at a site is not sufficient to estimate detectability

**Figure S4:**
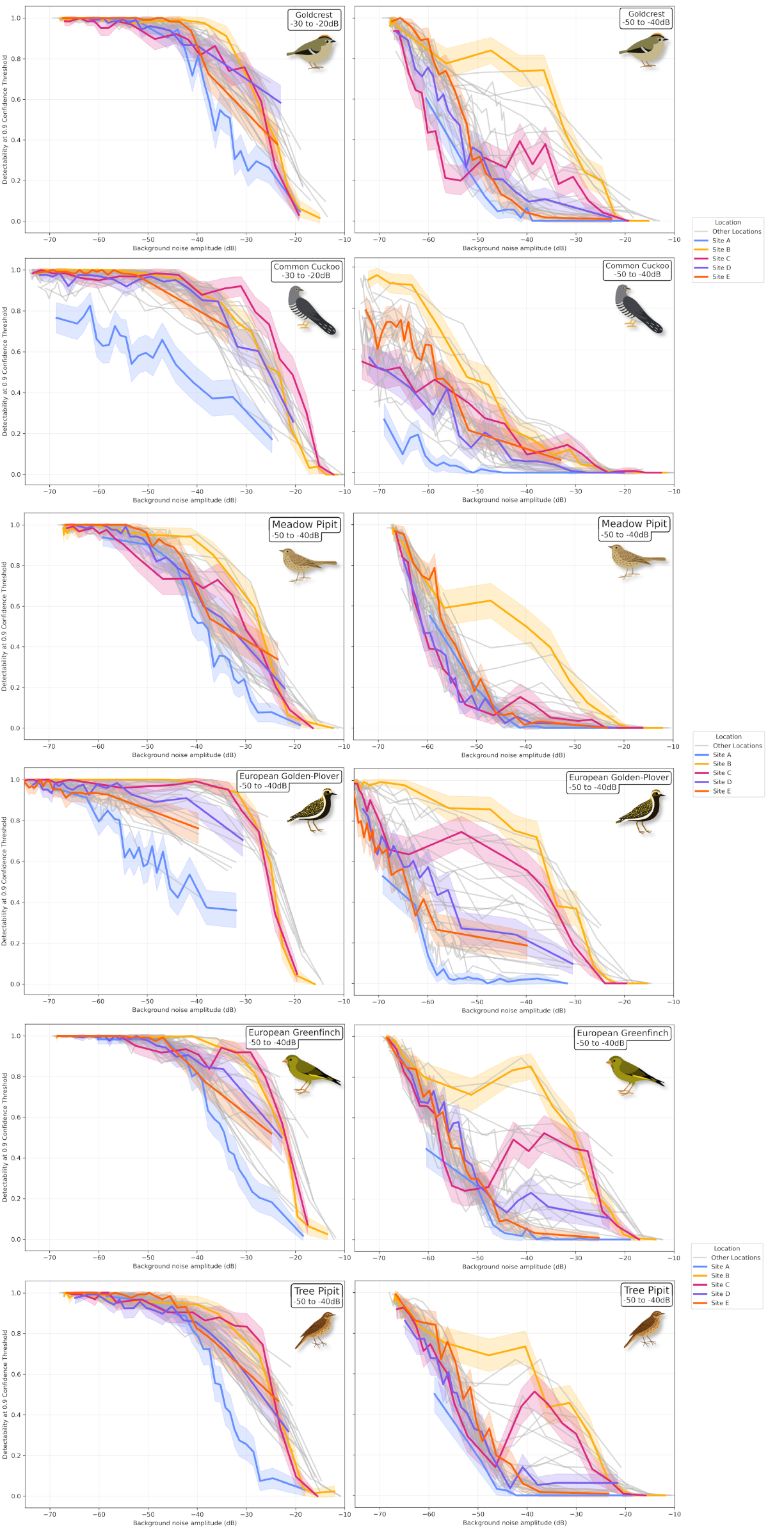
The relationship between background noise amplitude and detectability for all six species. Left column: high amplitude -30dB to -20dB vocalisations. Right column: average amplitude -50dB to -40dB vocalisations. Five of the 28 sites (A – E) are highlighted in colour.

## Footnotes

1 The specific version is: BirdNET-Analyzer v2.4.0, model BirdNET GLOBAL 6K V2.4 Model FP32.tflite

## References

Forrest, T. G. (1994). “From Sender to Receiver: Propagation and Environmental Effects on Acoustic Signals”. In: American Zoologist 34.6, pp. 644–654.

Gu, W. & R. K. Swihart (2004). “Absent or undetected? Effects of non-detection of species occurrence on wildlife–habitat models”. In: Biological Conservation 116.2. Publisher: Elsevier BV, pp. 195–203.

Slabbekoorn, H. (2004). “Habitat-dependent ambient noise: Consistent spectral profiles in two African forest types”. In: The Journal of the Acoustical Society of America 116.6, pp. 3727–3733.

Kéry, M. & B. Schmidt (2008). “Imperfect detection and its consequences for monitoring for conservation”. In: Community Ecology 9.2, pp. 207–216.

Bolker, B. M. et al. (2009). “Generalized linear mixed models: a practical guide for ecology and evolution”. In: Trends in Ecology & Evolution 24.3, pp. 127–135.

Simons, T. R., K. H. Pollock, J. M. Wettroth, M. W. Alldredge, K. Pacifici & J. Brewster (2009). “Sources of Measurement Error, Misclassification Error, and Bias in Auditory Avian Point Count Data”. In: Modeling Demographic Processes In Marked Populations. Ed. by D. L. Thomson, E.G. Cooch & M.J. Conroy. Boston, MA: Springer US, pp. 237–254.

Luther, D. & K. Gentry (2013). “Sources of background noise and their influence on vertebrate acoustic communication”. In: Behaviour 150.9-10, pp. 1045–1068.

Nemeth, E. et al. (2013). “Bird song and anthropogenic noise: vocal constraints may explain why birds sing higher-frequency songs in cities”. In: Proceedings of the Royal Society B: Biological Sciences 280.1754, p. 20122798.

Johnston, A. et al. (2014). “Species traits explain variation in detectability of UK birds”. In: Bird Study 61.3, pp. 340–350.

Buckland, S. T., E. A. Rexstad, T. A. Marques & C. S. Oedekoven (2015). Distance Sampling: Methods and Applications. Methods in Statistical Ecology. ISSN: 2199-319X, 2199-3203. Cham: Springer International Publishing.

Inger, R., R. Gregory, J. P. Duffy, I. Stott, P. Voříšek & K. J. Gaston (2015). “Common European birds are declining rapidly while less abundant species’ numbers are rising”. In: Ecology Letters 18.1. Ed. by J. Hill, pp. 28–36.

Darras, K., P. Pütz Fahrurrozi, K. Rembold & T. Tscharntke (2016). “Measuring sound detection spaces for acoustic animal sampling and monitoring”. In: Biological Conservation 201, pp. 29–37.

Brooks E. M., et al. (2017). “glmmTMB Balances Speed and Flexibility Among Packages for Zeroinflated Generalized Linear Mixed Modeling”. In: The R Journal 9.2, p. 378.

Guillera-Arroita, G. (2017). “Modelling of species distributions, range dynamics and communities under imperfect detection: advances, challenges and opportunities”. In: Ecography 40.2. Publisher: Wiley, pp. 281–295.

Yip, D. A., E. M. Bayne, P. Sólymos, J. Campbell & D. Proppe (2017). “Sound attenuation in forest and roadside environments: Implications for avian point-count surveys”. In: The Condor 119.1, pp. 73– 84.

Sebastián-González, E. et al. (2018). “Density estimation of sound-producing terrestrial animals using single automatic acoustic recorders and distance sampling”. In: Avian Conservation and Ecology 13.2, art7.

Baker, D. J. et al. (2019). “Conserving the abundance of nonthreatened species”. In: Conservation Biology 33.2, pp. 319–328.

Burns, P. A., C. McCall, K. C. Rowe, M. L. Parrott & B. L. Phillips (2019). “Accounting for detectability and abundance in survey design for a declining species”. In: Diversity and Distributions 25.10. Ed. By H. Regan. Publisher: Wiley, pp. 1655–1665.

Pérez-Granados, C., J. Gómez-Catasús, D. Bustillo-de La Rosa, A. Barrero, M. Reverter & J. Traba (2019). “Effort needed to accurately estimate Vocal Activity Rate index using acoustic monitoring: A case study with a dawn-time singing passerine”. In: Ecological Indicators 107, p. 105608.

Stowell, D., T. Petrusková, M. Šálek & P. Linhart (2019). “Automatic acoustic identification of individuals in multiple species: improving identification across recording conditions”. In: Journal of The Royal Society Interface 16.153, p. 20180940.

White, E. R. (2019). “Minimum Time Required to Detect Population Trends: The Need for Long-Term Monitoring Programs”. In: BioScience 69.1. Publisher: Oxford University Press (OUP), pp. 40–46.

Abrahams, C. & M. Geary (2020). “Combining bioacoustics and occupancy modelling for improved monitoring of rare breeding bird populations”. In: Ecological Indicators 112, p. 106131.

Devarajan, K., T. L. Morelli & S. Tenan (2020). “Multi-species occupancy models: review, roadmap, and recommendations”. In: Ecography 43.11. Publisher: Wiley, pp. 1612–1624.

Thomas, A., P. Speldewinde, J. D. Roberts, A. H. Burbidge & S. Comer (2020). “If a bird calls, will we detect it? Factors that can influence the detectability of calls on automated recording units in field conditions”. In: Emu Austral Ornithology 120.3, pp. 239–248.

Burns, F. et al. (2021). “Abundance decline in the avifauna of the European Union reveals cross-continental similarities in biodiversity change”. In: Ecology and Evolution 11.23, pp. 16647–16660.

Kahl, S., C. M. Wood, M. Eibl & H. Klinck (2021). “BirdNET: A deep learning solution for avian diversity monitoring”. In: Ecological Informatics 61, p. 101236.

Pérez-Granados, C. & J. Traba (2021). “Estimating bird density using passive acoustic monitoring: a review of methods and suggestions for further research”. In: Ibis 163.3, pp. 765–783.

Sánchez-Bayo, F. & K. A. G. Wyckhuys (2021). “Further evidence for a global decline of the entomofauna”. In: Austral Entomology 60.1, pp. 9–26.

Cole, J. S., N. L. Michel, S. A. Emerson & R. B. Siegel (2022). “Automated bird sound classifications of long-duration recordings produce occupancy model outputs similar to manually annotated data”. In: Ornithological Applications 124.2, duac003.

Leseberg, N. P., W. N. Venables, S. A. Murphy, N. A. Jackett & J. E. M. Watson (2022). “Accounting for both automated recording unit detection space and signal recognition performance in acoustic surveys: A protocol applied to the cryptic and critically endangered Night Parrot ( Pezoporus occidentalis )”. In: Austral Ecology 47.2, pp. 440–455.

Moussy, C. et al. (2022). “A quantitative global review of species population monitoring”. In: Conservation Biology 36.1. Publisher: Wiley.

Shaw, T., S. Müller & M. Scherer-Lorenzen (2022). “Slope does not affect autonomous recorder detection shape: considerations for acoustic monitoring in forested landscapes”. In: Bioacoustics 31.3, pp. 261–282.

Stowell, D. (2022). “Computational bioacoustics with deep learning: a review and roadmap”. In: PeerJ 10, e13152.

Waldock, C. et al. (2022). “A quantitative review of abundance-based species distribution models”. In: Ecography 2022.1. Publisher: Wiley.

Wood, C. M. & M. Z. Peery (2022). “What does ‘occupancy’ mean in passive acoustic surveys?” In: Ibis 164.4, pp. 1295–1300.

Brunk, K. M., R. J. Gutiérrez, M. Z. Peery, C. A. Cansler, S. Kahl & C. M. Wood (2023). “Quail on fire: changing fire regimes may benefit mountain quail in fire-adapted forests”. In: Fire Ecology 19.1, p. 19.

Ghani, B., T. Denton, S. Kahl & H. Klinck (2023). “Global birdsong embeddings enable superior transfer learning for bioacoustic classification”. In: Scientific Reports 13.1, p. 22876.

Haupert, S., F. Sèbe & J. Sueur (2023). “Physics-based model to predict the acoustic detection distance of terrestrial autonomous recording units over the diel cycle and across seasons: Insights from an Alpine and a Neotropical forest”. In: Methods in Ecology and Evolution 14.2, pp. 614–630.

Hutschenreiter, A., J.R. Sosa-López, F. González-García & F. Aureli (2023). “Evaluating factors affecting species detection using passive acoustic monitoring in neotropical forests: a playback experiment”. In: Bioacoustics 32.6, pp. 660–678.

Kelly, K. G. et al. (2023). “Estimating population size for California spotted owls and barred owls across the Sierra Nevada ecosystem with bioacoustics”. In: Ecological Indicators 154, p. 110851.

Pérez-Granados, C. (2023). “A First Assessment of Birdnet Performance at Varying Distances: A Playback Experiment”. In: Ardeola 70.2.

Ross, S. R. P. et al. (2023). “Passive acoustic monitoring provides a fresh perspective on fundamental ecological questions”. In: Functional Ecology 37.4, pp. 959–975.

Schmidt, B. R., S. S. Cruickshank, C. Bühler & A. Bergamini (2023). “Observers are a key source of detection heterogeneity and biased occupancy estimates in species monitoring”. In: Biological Conservation 283, p. 110102.

Sethi, S. S. et al. (2023). Automatic vocalisation detection delivers reliable, multi-faceted, and global avian biodiversity monitoring. preprint. Ecology.

Xie, J., Y. Zhong, J. Zhang, S. Liu, C. Ding & A. Triantafyllopoulos (2023). “A review of automatic recognition technology for bird vocalizations in the deep learning era”. In: Ecological Informatics 73, p. 101927.

Bennett, J. R. et al. (2024). “How ignoring detection probability hurts biodiversity conservation”. In: Frontiers in Ecology and the Environment 22.8. Publisher: Wiley.

Bielski, L., C. Cansler, K. McGinn, M. Z. Peery & C. Wood (2024). “Can the Hermit Warbler ( Setophaga occidentalis ) serve as an old-forest indicator species in the Sierra Nevada?” In: Journal of Field Ornithology 95.1, art4.

Callaghan, C. T., L. Santini, R. Spake & D. E. Bowler (2024). “Population abundance estimates in conservation and biodiversity research”. In: Trends in Ecology & Evolution 39.6, pp. 515–523.

Hutschenreiter, A. et al. (2024). “How to count bird calls? Vocal activity indices may provide different insights into bird abundance and behaviour depending on species traits”. In: Methods in Ecology and Evolution 15.6, pp. 1071–1083.

Jarrett, D. & S. G. Willis (2024). “Acoustic detection rate can outperform traditional survey approaches in estimating relative densities of breeding waders”. In: Ibis, ibi.13375.

Lebeuf-Taylor, I., E. Knight & E. Bayne (2024). “Improving bird abundance estimates in harvested forests with retention by limiting detection radius through sound truncation”. In: Ornithological Applications, duae055.

Navine, A. K., R. J. Camp, M. J. Weldy, T. Denton & P. J. Hart (2024). “Counting the chorus: A bioacoustic indicator of population density”. In: Ecological Indicators 169, p. 112930.

Serrurier, A. et al. (2024). “Mountain is calling – decrypting the vocal phenology of an alpine bird species using passive acoustic monitoring”. In: Ibis, ibi.13314.

Wernberg, T. et al. (2024). “Impacts of Climate Change on Marine Foundation Species”. In: Annual Review of Marine Science 16.1, pp. 247–282.

Winiarska, D., P. Szymański & T. S. Osiejuk (2024). “Detection ranges of forest bird vocalisations: guidelines for passive acoustic monitoring”. In: Scientific Reports 14.1, p. 894.

Jarrett, D. et al. (2025). “Mitigating bias in long-term terrestrial ecoacoustic studies”. In: Journal of Applied Ecology, pp. 1365–2664.70000.

Lindner, K. et al. (2025). “Sampling effort for community composition higher by a magnitude compared to species richness in passive acoustic monitoring”. In: Ecological Informatics 90, p. 103265.

Massimino, D. et al. (2025). “The Breeding Bird Survey of the United Kingdom”. In: Global Ecology and Biogeography 34.1, e13943.

Xeno-canto Foundation (2025). Xeno-canto: Sharing wildlife sounds from around the world.

